# Evolution of body size and wing shape trade-offs in arsenurine silkmoths

**DOI:** 10.1101/2020.05.12.092197

**Authors:** Chris A. Hamilton, Nathalie Winiger, Juliette J. Rubin, Jesse Breinholt, Rodolphe Rougerie, Ian J. Kitching, Jesse R. Barber, Akito Y. Kawahara

## Abstract

One of the key objectives in biological research is understanding how evolutionary processes have produced Earth’s biodiversity. These processes have led to a vast diversity of wing shapes in insects; an unanswered question especially pronounced in moths. As one of the major predators of nocturnal moths, bats are thought to have been involved in a long evolutionary arms race with their prey. In response, moths are thought to have evolved many counter strategies, such as diverse wing shapes and large body sizes. However, the tradeoffs between body size and wing shape are not well understood. Here we examined the evolution of wing shape in the wild silkmoth subfamily Arsenurinae (Saturniidae). By using phylogenomics and geometric morphometrics, we established the framework to evaluate potential evolutionary relationships between body size and wing shape. The phylogeny was inferred based on 781 loci from target capture data of 42 arsenurine species representing all 10 recognized genera.

We found there are evolutionary trade-offs between body size, wing shape, and the interaction of fore- and hindwing shape. Namely, body size decreases with increasing hindwing length, but increases as forewing shape becomes more complex. Additionally, hindwing shape has a significant effect on forewing shape complexity. The complex wing shapes that make Arsenurinae, and silkmoths as a whole, so charismatic are likely driven by the strong forces of natural selection and genomic constraints.

One other important outcome was discovering within our data one of the most vexing problems in phylogenetic inference – a region of a tree that possesses short branches and no “support” for relationships (i.e., a polytomy). These parts of the Tree of Life are often some of the most interesting from an evolutionary standpoint. To investigate this problem, we used reciprocal illumination to determine the most probable generic relationships within the Arsenurinae by inspecting differing phylogenetic inferences, alternative support values, quartets, and phylogenetic networks to reveal hidden phylogenetic signal.

---

The vast amount of Earth’s diversity in faunal form and function lies in the arthropod Tree of Life, yet major questions persist: 1) How many arthropod species wait to be discovered and described? 2) What are the relationships across the arthropod Tree of Life, particularly towards the tips? and 3) What characters, traits, or interactions have allowed some lineages to become more diverse than others? The lineage that provides perhaps the most informative opportunities for answering these questions is the Insecta. Arguably the most successful lineage on the planet, insects have diversified to fill virtually all terrestrial and freshwater niches ((Ehrlich and Raven 1964), (Farrell et al. 1992), (Mitter et al. 1988), (Wiens et al. 2015), (Condamine 2016)), often evolving incredible traits to exploit them, such as wings and the ability to fly. These innovations provided both ecological opportunities and challenges to diversify in the face of new pressures, particularly predatory.

Between 50-70 million years ago, a major lineage of insect predator appeared - echolocating bats ((Jones et al. 2005), (Teeling et al. 2005), (Miller-Butterworth et al. 2007), (Shi and Rabosky 2015), (Lei and Dong 2016)). Nightly battles between moths and bats drove a predator-prey arms race ((Corcoran et al. 2009), (Conner and Corcoran 2012)) that produced remarkable anti-bat strategies such as ultrasonic-detecting ears ((Roeder and Treat 1957), (Scoble 1992)), ultrasound producing organs capable of jamming bat sonar ((Corcoran et al. 2011), (Barber and Kawahara 2013)) or warning of bad taste ((Dunning 1968), (Hristov and Conner 2005), (Barber et al. 2009), (Dowdy and Conner 2016)), and evasive flight strategies – such as aerial maneuvers or temporal partitioning of activity ((Lamarre et al. 2015), (Kawahara et al. 2018)). Amidst the emergence of these fierce predators, one of the most captivating lepidopteran radiations arose – the superfamily Bombycoidea ((Wahlberg et al. 2013), (Misof et al. 2014), (Kawahara and Barber 2015), (Kawahara et al. 2019)).

Some of the most spectacular anti-bat strategies can be found in the bombycoid sister lineages Saturniidae (wild silkmoths) and Sphingidae (hawkmoths) – two lineages with an incredible array of shapes and behavioral traits ((Barber and Kawahara 2013), (Breinholt and Kawahara 2013), (Kawahara and Breinholt 2014), (Barber et al. 2015), (Rubin et al. 2018)). The divergent life-history strategies of these two lineages has likely played a major role in driving their diversity (Hamilton et al. 2019). For example, the majority of hawkmoths are “income breeders” ((Janzen 1984), (Tammaru and Haukioja 1996)); adults live for a relatively long time period (weeks to months) during which they feed on nectar while traveling long distances looking for mates, mate multiple times, retain the eggs internally for long periods of time to allow egg maturation, and searching for their appropriate larval host plants (frequently highly specific and toxic) upon which to lay their eggs. Their incredibly fast and maneuverable flight, including the ability to hover and fly backwards, as well as their sleek appearances, has earned them a popular caricature as the “fighter jets” of the moth world ((Roeder 1974), (Rydell and Lancaster 2000)). In addition, many sphingid lineages possess ears or ultrasound producing organs that have independently evolved multiple times to detect and respond to their echolocating predators ((Barber and Kawahara 2013), (Kawahara and Barber 2015)), whereas some sphingids possess neither. This apparent vulnerability begs the question: How do lineages that cannot hear bat echolocation survive the nightly gauntlet?

Initial answers can be found in the Saturniidae, generally the most charismatic lineage of moth due to their large body sizes, striking colors and patterns, and elaborate wing shapes. As “capital breeders”, they possess a very different life-history strategy from their sister lineage ((Janzen 1984), (Tammaru and Haukioja 1996)). Adults typically only live one or two weeks after emergence, and importantly, they do not feed as adults as they lack functional mouthparts (see Janzen 1984). Most of the saturniid life span encompasses the larval period, during which they do not sequester toxins – a markedly different strategy from their sister lineage. It is during this critical stage that they gather as many resources as possible to fuel the adult stage, where they usually mate once with the first available mate and the female almost immediately lays her eggs on what are generally non-toxic (but nutritionally poor) plants (e.g., most saturniids feed on trees, whereas most sphingids feed on flowering plants) ((Scoble 1992), (Morton 2009)). As adults, saturniids possess neither ears nor ultrasound-producing organs, yet they are far from defenseless. As with the sphingids, saturniids are often quite large moths, and this sheer body size itself could forestall bat attack ((Roeder 1974), (Jacobs and Bastian 2016); also see Discussion). Saturniid bodies are also covered in dense scales, creating a “furry” appearance, that absorb bat biosonar and may effectively reduce the moth’s detectability distance by bats (Neil et al. 2020). Once detected, some saturniids thwart their echolocating bat prey by luring attacks to non-essential wing areas. A number of species possess long hindwing tails with twisted and cupped ends that rotate behind them in flight, likely creating a sensory illusion that fools the bat into attacking these appendages ((Barber et al. 2015), (Rubin et al. 2018)). The efficacy of this trait is scaled with hindwing tail length, as increasingly long tails increasingly draw bat attack towards these non-essential appendages, away from the vital body core, allowing the moth to escape more often (Rubin et al. 2018). While long tails provide the greatest benefit for predator escape, simply having an elongated hindwing also provides protection; experimentally elongating the hindwing of a saturniid moth elevates its escape success ∼25% (Rubin et al. 2018). Interestingly, studies on multiple Lepidoptera groups have shown that severe damage or entire removal of the hindwings does not greatly inhibit flight, while similar forewing damage or removal renders the animal flightless ((Jantzen and Eisner 2008), (Le Roy et al. 2019), (Stylman et al. 2019)). Within the Saturniidae, several subfamilies and multiple genera include tailed and non-tailed species, providing an excellent opportunity to understand the evolution of lineages with anti-bat wing traits and those without.

One of these subfamilies, the Arsenurinae, are large, cryptic moths that mostly inhabit low to mid-elevation tropical forests throughout the Neotropics (Lemaire and Minet 1998). Importantly, across the 10 genera and 89 species (Kitching et al. 2018), they possess a tremendous amount of divergence in wing shape ((Zhong et al. 2016), (Barber et al. 2015), (Rubin et al. 2018)). Phylogenetic relationships among the genera of Arsenurinae have been hypothesized three times ((Michener 1952), (Peigler 1993), (De Camargo et al. 2009)), with major differences in the generic relationships postulated each time (Supp. Fig. 1). While the study of Michener (1952) was pre-cladistic, it contrasted derived and ancestral character states as indicators of phylogenetic affinities. Peigler (1993) and DeCamargo et al. (2009) undertook cladistic studies that used morphological characters to infer the monophyly of the Arsenurinae and its constituent genera. Unfortunately, the only consistent phylogenetic relationships on which they all agreed were that *Dysdaemonia* and *Titaea* were sister genera, and that *Almeidaia* was sister to all other Arsenurinae (Supp. Fig. 1). As a result, we have lacked a robust phylogeny and understanding of relationships that could be used to test hypotheses regarding the group’s evolution. Recently, Rubin and Hamilton et al. (2018) densely sampled the Saturniinae, using phylogenomics, to provide the clearest picture so far of the relationships within this subfamily; however, that study lacked sampling outside of the Saturniinae. Hamilton et al. (2019) then examined relationships within the superfamily Bombycoidea and identified diversification rate shifts that may be attributed to the predatory pressure of bats, but Arsenurinae sampling was severely limited with only two lineages included.

To build the evolutionary framework from which we can trace the path of potential anti-bat traits, we inferred the first molecular phylogeny of the subfamily Arsenurinae. We subsequently investigated the evolution of wing shape and body size by using geometric morphometrics from natural history collection specimens and tested whether there have been evolutionary trade-offs between body size, fore- and hindwing shape, and the presence of hindwing tails as a hypothesis-generating approach for future trait-based work.

## Material and Methods

### Taxon Sampling

We sampled 36 out of 89 described species from the ten arsenurine genera (see Kitching et al. 2018): *Almeidaia* Travassos, 1937; *Arsenura* Duncan, 1841; *Caio* Travassos & Noronha, 1968; *Copiopteryx* Duncan, 1841; *Dysdaemonia* Hübner, [1819]; *Grammopelta* Rothschild, 1907; *Loxolomia* Maassen, 1869; *Paradaemonia* Bouvier, 1925; *Rhescyntis* Hübner, [1819]; and *Titaea* Hübner, [1823]. To examine the placement of Arsenurinae wing shape morphospace within the context of the Saturniidae, we digitally imaged and quantified the shape complexity (as principal components) of 968 male specimens comprising 174 species across the eight subfamilies (see Supp. Tables 2 & 3). Ingroup sampling of images (i.e., the Arsenurinae) comprised 545 specimens corresponding to 50 species of the 89 total ((Kitching et al. 2018); see Supp. Tables 4 & 5). Outgroup lineages were chosen because they possess high variation in wing shape morphology, allowing the capture of the morphological diversity outside of the Arsenurinae.

### Molecular Data

We obtained specimens for phylogenetic analyses from the molecular collections of the McGuire Center for Lepidoptera and Biodiversity (MGCL) at the Florida Museum of Natural History (FLMNH), the University of Maryland (UMD), and the Muséum national d’Histoire naturelle in Paris (MNHN). Twenty-six pinned or papered museum specimens, some as old as 36 years, were used for sequencing, to compare recovery of DNA with 16 traditionally-stored molecular collections specimens in ≥95% ethanol and -80°C (see (McGaughran 2020); Supp. Table 1, Supp. Texts 1 & 2).

Genomic DNA from molecular collection specimens was extracted from the thorax or leg(s) using OmniPrep Genomic DNA Extraction Kits (G-Biosciences, St. Louis, MO, USA) or DNeasy Blood and Tissue Kits (Qiagen, Valencia, CA, USA). We used the protocol for extracting DNA from historical museum specimens that was developed and outlined in Hamilton et al. (2019). DNA concentration was evaluated through agarose gel electrophoresis and fluorometry using a Qubit 2.0 (Invitrogen, Thermo Fisher Scientific). The LEP1 Anchored Hybrid Enrichment (AHE) targeted sequencing kit (Breinholt et al. 2018), an Agilent Custom SureSelect Target Enrichment kit, was used to target 855 loci. Library preparation and Illumina HiSeq 2500 sequencing (PE100) was carried out at RAPiD Genomics, Gainesville, FL, USA. Tissues from the molecular samples, as well as all extracts, are preserved at -80C in the MGCL; wing vouchers are stored following the methods of Cho et al. (2016) and stored at the MGCL.

A previously developed bioinformatics pipeline was used to prepare sequences for phylogenetic inference (Breinholt et al. 2018). The pipeline uses a probe-baited iterative assembly that extends beyond the probe region, checks for quality and cross contamination due to barcode leakage, removes paralogs, and returns a set of aligned orthologs for each locus and taxon of interest. To accomplish these tasks, the pipeline uses the *Bombyx mori* genome (Xia et al. 2004) and the LEP1 AHE reference library. Loci for phylogenetic analysis were selected by applying a cutoff of ≥50% sampled taxa recovery (i.e., for a locus to be included in the analysis, the locus had to be recovered in at least 50% of the sampled taxa).

The pipeline evaluates density and entropy at each site of a nucleotide sequence alignment. As established by Hamilton et al. (2019), we elected to trim with entropy and density cutoffs only in “flanking” regions, allowing the “probe” region to be converted into amino acid sequences. For a site (outside of the probe region) to remain, that site must pass a 60% density and 1.5 entropy cutoff, rejecting sites that fail these requirements. A higher first value (60) increases the coverage cutoff (e.g., a site is kept if 60% of all taxa are represented at that site). A higher second value (1.5) increases the entropy cutoff (i.e., entropy values represent the amount of saturation at a site); sites with values higher than 1.5 possess higher saturation and are thus deleted. AliView v1.18 (Larsson 2014) was used to translate to amino acids, check for frame shifts, recognize and remove stop codons (if present), and edit sequencing errors or lone/dubious indels. Because flanking sequences are generally non-coding and sites have been deemed homologous (following assembly and alignment), these flanking sequences were separated from the exons, then combined and treated together as an independent partition. Following the filtering steps in the bioinformatics pipeline (i.e., site orthology, and density and saturation evaluation), the flanking partition is viewed as a SNP supermatrix, where each site is homologous, but uninformative sites, saturated sites, or sites with large amounts missing data have been removed. All pipeline analyses, including phylogenomic analyses (below) were conducted on the University of Florida High-Performance Computing Center HiPerGator2 (http://www.hpc.ufl.edu/).

### Phylogenetics

Phylogenomic inferences were conducted under likelihood optimality criteria in IQ-TREE multicore version 1.6.9 (Nguyen et al. 2015). To examine phylogenetic signal, we evaluated nucleotide and amino acid datasets. Four supermatrix datasets were built for phylogeny inference: 1) AA = an amino acid supermatrix (777 loci) composed of translated probe region loci; 2) Pr = probe region-only, as nucleotides (778 loci; each locus modeled by sites); 3) Pr+Fl = a probe + flanking supermatrix (782 loci; each locus modeled by sites); and 4) Fl = flanking supermatrix only (modeled by sites). Because concatenation has been shown to fail when there are high levels of incomplete lineage sorting (see Mendes and Hahn 2017), we assessed the impact of potential gene-tree discordance ((Maddison 1997), (Slowinski and Page 1999), (Edwards 2009)) by inferring a phylogeny for each individual locus from the Pr+Fl supermatrix using IQ-TREE, with subsequent species tree estimation performed in ASTRAL-III version 5.6.3 ((Mirarab et al. 2014), (Mirarab and Warnow 2015), (Zhang et al. 2017)) (see Supplemental Trees). For both nucleotide and amino acid datasets, the ‘–m TEST’ command was used in IQ-TREE to perform a search for the most appropriate model of amino acid or nucleotide substitution. Traditional branch support values were computed via 1000 random addition sequence (RAS) replicates, and 1000 replicates each for both ultrafast bootstraps (UFBS) (‘–bb’ command), as well as SH-aLRT tests (‘-alrt’ command). Nodes were classified as “robust” if they were recovered with support values of UFBS ≥ 95 and SH-aLRT ≥ 80 ((Minh et al. 2013), (Guindon et al. 2010)). ASTRAL support values (ASV) – local posterior probabilities (Sayyari and Mirarab 2016), were used to evaluate node support on the species tree. ASTRAL support values were determined to be “robust” if nodes were recovered with local posterior probabilities ≥ 0.95.

Because phylogenomic datasets can produce incongruent topologies and artificially inflate traditional nodal support values (e.g., due to the increase in total number of sites, the sampling variance for a particular branch is low - see Reddy et al. 2017), we evaluated additional modes of topological support. To identify potential rogue taxa or unstable regions in the tree, we ran RogueNaRok ((Aberer and Stamatakis 2011), (Aberer et al. 2012)), http://rnr.h-its.org/, on the 1000 ultrafast bootstrap trees and the consensus tree from the Pr+Fl supermatrix. Potential rogue taxa were removed, and phylogenetic inference was rerun (‘AA minus *Grammopelta*, ‘Pr+Fl minus *Grammopelta*’). Disagreements among loci were estimated using concordance factors (gCF – gene concordance factor of (Minh et al. 2018)) calculated in IQ-TREE 1.7-beta12, on both the Pr+Fl supermatrix constrained tree #1 (see below and Supplemental Trees) and the ASTRAL tree. Internode Certainty scores (IC) were calculated using QuartetScores (Zhou et al. 2020) to infer a measure of support for the reference topology compared with the frequency of the most prevalent alternative topology. Quartets were then investigated to inspect relationships at nodes with low support and evaluate whether one topology was occurring more often than others. Quartet testing was carried out using Likelihood Mapping (Strimmer and vonHaeseler 1997) in IQ-TREE (Nguyen et al. 2015). In this hypothesis-testing framework, we computed the likelihood of possible relationships that can be constructed from possible quartets of the taxa. For example, we looked at the subfamily level by testing whether Arsenurinae was more closely related to the Agliinae, Salassinae, or Saturniinae. We also investigated the genus level by grouping lineages, according to the phylogeny, and evaluating (from a quartet standpoint) whether lineages were found more closely related to another. Quartet likelihood mapping is also a way to inspect the information (i.e. phylogenetic) content of a dataset. To investigate potential gene tree/species tree incongruence, we computed a phylogenetic network in SplitsTree4 version 4.14.4 (Huson and Bryant 2006). Because the gene trees do not include identical sets of taxa, the ‘SuperNetwork’ method was implemented, using uncorrected p-distance and Jukes-Cantor models, in which the edge weights equaled “TreeSizeWeightedMean”. We attempted to investigate networks using PhyloNet 3 ((Than et al. 2008), (Wen et al. 2018)). Due to the computational resources this approach requires, we reduced the taxon set to a single representative for each genus and attempted to evaluate the “best” total log probability score from the inferred networks, after testing whether the number of reticulations had been from 0 to 14 (the total number of tips in this reduced taxon set). Finally, using the process of reciprocal illumination, we coalesced these outcomes and reran phylogenetic inference with two unrooted constrained topologies: (#1) a simplified tree with the major generic relationships reduced to a polytomy ((((*Copiopteryx, Rhescyntis*),(*Titaea, Dysdaemonia*), *Grammopelta, Paradaemonia, Caio, Loxolomia*), *Arsenura*), *Almeidaia*, (*Salassa, Aglia*)); and (#2) a simplified tree based on the topology of the ‘Pr+Fl minus *Grammopelta’* to see where *Grammopelta* would be placed (see Supplemental Trees). To test whether one topology was better than another, we ran Shimodaira-Hasegawa tests in the R package ‘phangorn’ (Schliep 2010), under 10,000 bootstrap replicates. All trees were rooted with six outgroup species from the subfamilies Saturniinae, Salassinae, and Agliinae (Supp. Table 1). All consensus alignment FASTA files, loci information, partition files, tree files, and other essential data files used for phylogenetic inference are available as supplementary materials on the Dryad Data Repository (doi:???). All data matrices and resulting trees are available in TreeBASE (???).

### Digitization

The majority of specimens digitized were from the MGCL collection. Additional images of *Almeidaia aidae, Caio richardsoni*, and *Arsenura drucei* were obtained from the Barcode of Life Datasystems (BOLD); www.boldsystems.org (Ratnasingham and Hebert 2007). Further images of *Almeidaia aidae* were obtained from specimens at the MNHN and from Eurides Furtado (Furtado 2004) (see the non-LEP/non-MGCL codes in Supp. Tables 2-5). Specimens from these alternative sources were used because these species were unavailable in the MGCL (e.g., *Almeidaia* specimens are exceedingly rare in collections). Images of the right wing (or left wing, if right side was damaged or broken) were taken with a Canon EOS Rebel T3i digital camera with EF 35mm F2.0 lens. Pinned specimens were mounted onto a piece of clay and carefully placed on an opaque acrylic sheet (355 × 355 x 3 mm) in a way that the wings were in a horizontal plane. If needed, the wings of pinned specimens were slightly moistened with 90% EtoH in order to provide transparency for visualizing the area of overlap between the forewing (FW) and hindwing (HW). A metric ruler was included with each image to provide scale. If the fore- and hindwing were not in one plane, two images were taken.

Measurements used for comparative analyses included FW length (FW_L), HW length (HW_L), and type of HW tail shape as a categorical variable. FW length was chosen because it is widely used as a proxy for body size (Miller 1977). Images were processed in ImageJ (Abràmoff et al. 2004) to obtain measurements of the wings. FW and HW length were measured using the included ruler for scale. Forewing length was measured as the straight-line distance between a point placed at the middle of the base of the wing junction to the thorax, and one placed at the apex of the forewing (Supp. Fig. 2a). Hindwing length was measured from a point placed at the middle of the wing junction to the thorax, to the point of the deepest curve (convex) along the tornal edge of the hindwing (i.e., most prominent extension or curvature on the tornal margin) or the tornal end of the hindwing tail (Supp. Fig. 2b). Photoshop CS6 Version 13.0 was used to obtain the wing shape outlines used in the Elliptical Fourier analyses. To do this, FW and HW were highlighted and cut out separately with the quick selection tool (standard set up: size 22 pixels, hardness 100%, and spacing 25%). The wing of choice was selected and colored pure black, with the background converted to pure white. Images were converted to black and white with the greyscale function. If images of the left wings were used, they were flipped horizontally to have the same orientation as the other images. Specimens imaged at the MGCL had a green label with “MGCL ####” added with the specimen. All images are available in the supplemental information within the Dryad Digital Repository (doi:???).

### Elliptical Fourier Descriptor analysis

Analyzing shape variation among objects can reveal insights into their function and the underlying mechanisms leading to their variation. Geometric morphometrics (GM) is a powerful tool to study phenotypic variation and covariation, as size and shape are treated as separate variables (Zelditch et al. 2012). Elliptical Fourier Descriptor (EFD) methods were developed for fitting curves to complex closed contours (Kuhl and Giardina 1982) and can be used to analyze the outlines of objects and numerically describe shapes that have few or no identifiable homologous (i.e., explicit) landmarks ((Iwata et al. 1998), (Chitwood 2014), (Bonhomme et al. 2013), (Bonhomme et al. 2014)), while eliminating size as a variable in shape. Instead of using traditional landmark data to quantify Lepidoptera wing shape (e.g., (Nath and Devi 2009), (Chazot et al. 2016), (Zhong et al. 2016)), we investigated the usefulness of EFD using the R program ‘Momocs’ (Bonhomme et al. 2014). EFD has been used frequently in plants to study the evolution of leaf shape ((Chitwood and Naylor 2012), (Chitwood et al. 2012a), (Chitwood et al. 2012b), (Chitwood et al. 2013), (Chitwood et al. 2014)), but rarely in animals ((Rohlf and Archie 1984), (Felice and O’Connor 2014), (Zhan and Wang 2012), (Sharma et al. 2017)). Recently however, it has been shown to be a powerful approach to study shape evolution in Lepidoptera ((Rubin et al. 2018), (Hegedus et al. 2018)). We chose male specimens because they fly long distances looking for mates, while females generally sit in the vegetation very near where they emerge from the pupa (e.g., Arsenurinae pupate in the soil), waiting for males to find them, which thus potentially exposes males to higher bat predation pressure (see (Rutowski 1982), (Acharya 1995)).

To quantify wing shape and morphospace, the first harmonic (i.e., defines the best-fitting ellipse) was used to normalize the harmonic coefficients and render them invariant to size and rotation. This approach allows quantifiable analysis of shape when there are few or no identifiable homologous landmarks ((Iwata and Ukai 2002), (Bonhomme et al. 2014), (Chitwood 2014), (Chitwood and Sinha 2016)). EFD uses the first ellipse to normalize for rotation, translation, size and orientation, then uses harmonic coefficients for subsequent statistical analysis, Principle Components Analysis (PCA), and visualization. Outlines were defined as the closed polygon formed by the (x; y) coordinates of bounding pixels. A smoothing iteration was visually determined by eye and set to 500 for both HW and FW, using the function ‘coo_smooth’. The harmonics were then estimated using the function ‘calibrate_harmonicpower_efourier’ to evaluate the number needed to effectively describe the shapes in this analysis, without overparameterization, and where 99% of the power is captured. For forewings (FW) the number of harmonics were set at 8, while the hindwings (HW) were set at 9. Prior to EFD, all images were stacked and centered on top of each other (‘stack’ and ‘coo_center’), scaled to the same size (‘coo_scale’), and then put into the same direction (‘coo_slidedirection’) to avoid poor alignment and rotation of the shapes. The harmonic coefficients were then used for subsequent statistical analysis (EFD), PCA to reduce the dimensionality of the efourier coefficients, and visualization of the morphospace. HW morphospace was then plotted and visualized for all specimens designated by subfamily, tribe, genus, or species, and HW shape category (i.e., “none” = no tail, “lobe” = bulge of HW material (both small and long), “small” tail, “medium” tail, “long” tail, and “extra long” tail), as well as for the Arsenurinae (ingroup) using only the genus, species, and HW shape category descriptor. FW morphospace was plotted and visualized using the same categories, in particular the HW shape category, in order to visualize potential relationships between the two variables.

### Trait Evolution

Phylogenetic relationships present problems for statistical inference because lineages are evolutionarily related and therefore the data are not independent. Our major questions asked whether there were evolutionary trade-offs between body size, wing shape, and the presence of hindwing tails. To evaluate the appropriateness of testing for statistical significance, we investigated the phylogenetic signal (Pagel’s λ) in our measured traits (i.e., HW shape, FW shape, body size (FW length = proxy for body size in moths), and HW length). After accounting for phylogeny, we tested for the effects of one trait on another by performing Phylogenetic Generalized Least Squares (PGLS) regression, under the ‘mvOU’ (multivariate Ornstein-Uhlenbeck) model of evolution (i.e., the mvOU model provided a better fit to the data than the BM (Brownian Motion) or EB (Early Burst) models) using the R package ‘Rphylopars’ (Goolsby 2017a); this program can estimate evolutionary covariance while accounting for within-species variation and missing data (see (Bruggeman et al. 2009), (Goolsby 2015), (Goolsby 2017b), (Goolsby 2017a)). In order to probe our dataset for possible evolutionary tradeoffs, we asked these questions: 1. Is HW length a predictor of FW_L [body size]?; 2. Is FW_L [body size] a predictor of HW length?; 3. Is HW length a predictor of FW shape complexity?; and 4. Is FW shape complexity a predictor of HW length? (see Table 1). Additionally, phylogenetic ANOVA were conducted using the R package ‘phytools’ (Revell 2011) to test whether the type of hindwing (as multistate categorical variables) had an effect on another trait (HW shape, FW shape, or body size). We coded the HW categories into discrete states (based on the results of Rubin et al. 2018): none (i.e., no HW tail) = 0, a small lobe = 1, a long lobe = 2, a small tail = 3, medium tail = 4, or extra long tail = 5 (Fig. 2). To do this, we reduced our dataset to only our ingroup, the Arsenurinae, by removing the outgroup tips from our preferred tree, the constrained tree (#1).

**Table 1.** Trait tests between body size (measured as FW length - a proxy for body size), and FW shape and HW length (as principal components). Information included are the specific tests and their significance.

| Questions | PGLS regression | significance? | p-value |
| --- | --- | --- | --- |
| Is HW length a predictor of FW_L [body size]? | FW_L ~ HW_L | YES | 0.001115 |
| Is FW_L [body size] a predictor of HW length? | HW_L ~ FW_L | NO | 0.1131 |
| Is HW length a predictor of FW shape complexity? | FW_PC1 ~ HW_L | YES | 2.20E-16 |
| Is FW shape complexity a predictor of HW length? | HW_L ~ FW_PC1 | NO | 0.8423 |

**Figure 1.**
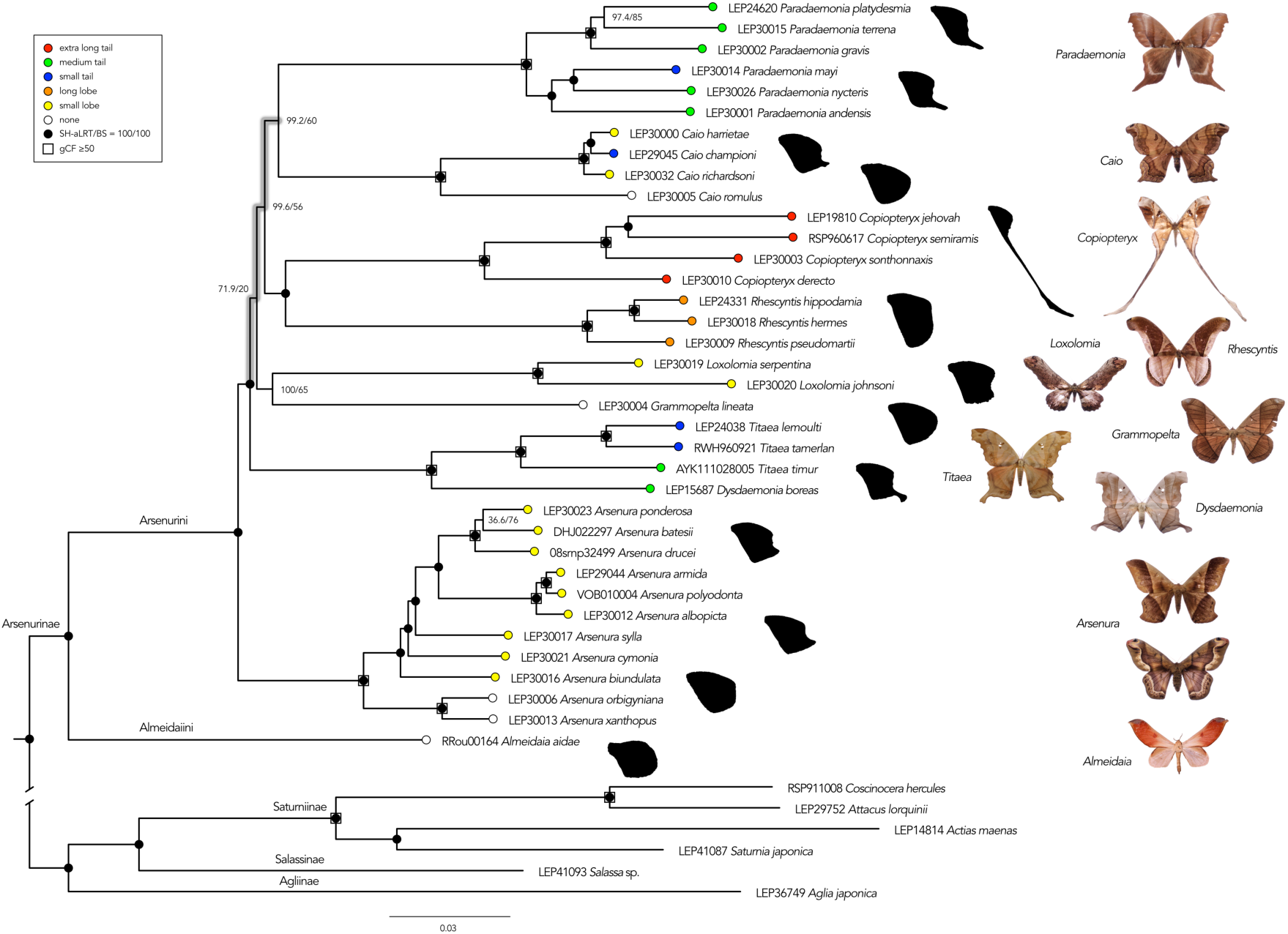
The preferred Arsenurinae phylogeny, constrained inference (#1), based on 782 AHE loci. Grey shading on branches indicate the problematic region along the backbone of the phylogeny. Nodes with support values with SH-aLRT & bootstrap (BS) values = 100% are indicated with a black circle. Support values below these are stated. Nodes with concordance factors ≥ 50 are indicated with a white box. Hindwing tail designations: extra-long tail (red), medium tail (green), small tail (blue), long lobe (orange), small lobe (yellow), no tail (white). Generalized wing shapes for genera or species groups within a genus are placed at the tips, in black. Photographs represent a generalized species from the genera sampled in the phylogeny.

**Figure 2.**
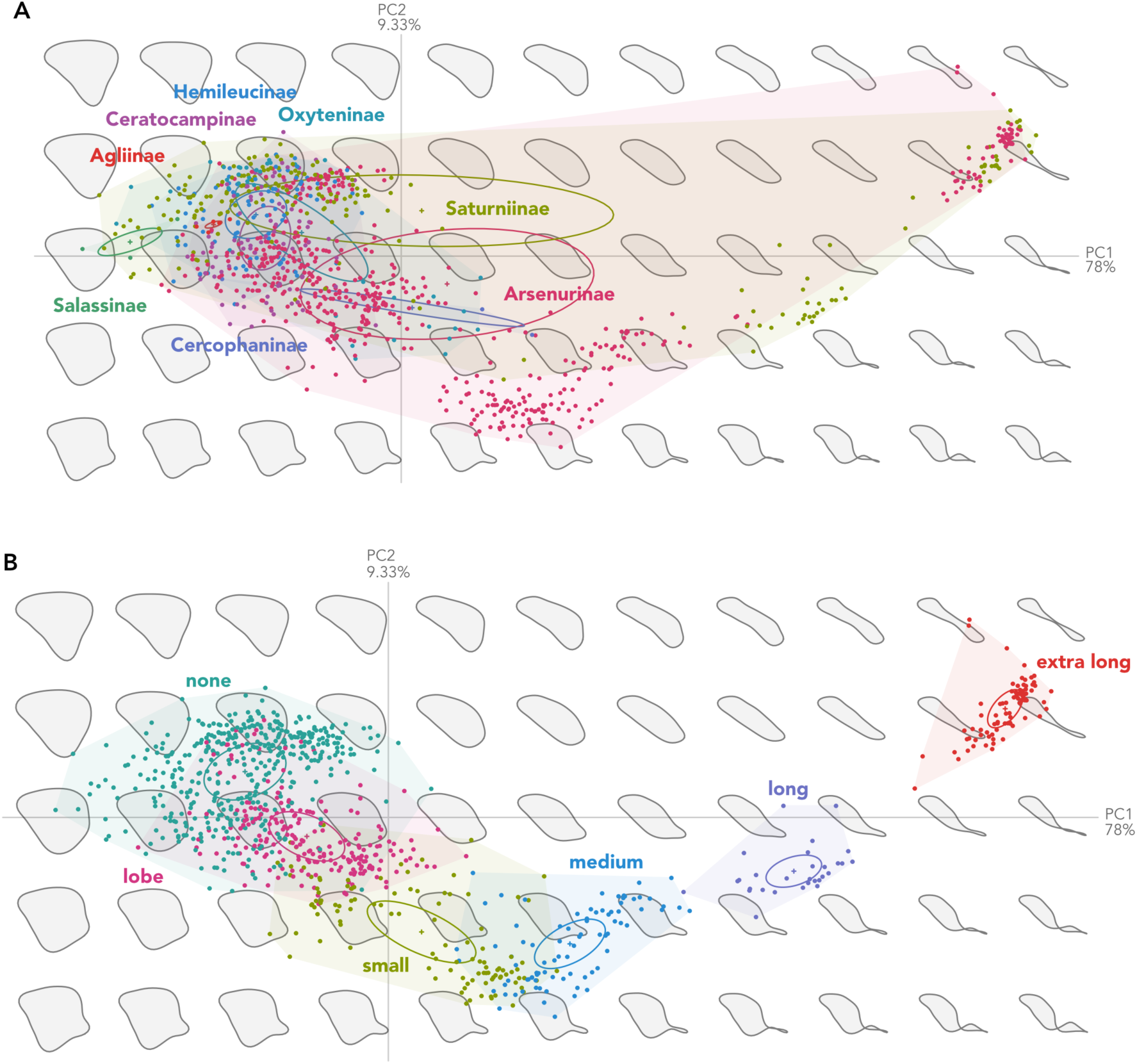
Geometric morphometric analysis of hindwing (HW) shape reveals changes in morphospace across the Saturniidae subfamilies. Principal components (PC1∼PC2) are plotted to visualize hindwing morphospace by A) subfamily, showing Saturniinae and Arsenurinae have convergently evolved into the same morphospace; and B) Hindwing (HW) shape morphospace plotting the hindwing tail categories “none”, “lobe”, “small”, “medium”, “long”, and “extra long”. Dots represent individual specimens in the analysis. Hypothetical shape approximations are plotted in the background to aid in visualizing shape change.

The tree was then converted into a relative-rate scaled ultrametric tree using the ‘chronopl’ command in the R package ‘ape’ (Paradis and Schliep 2019). This approach produces a tree whose branches are scaled to evolutionary rates, rather than a dated tree, and provides a means to understand evolutionary changes over relative “time” in the group being investigated. Our quantitative dataset was then matched to the ingroup species and tips in the tree. Trait distributions were also visualized using the contMap function in ‘phytools’ (Revell 2011).

## Results

### Phylogenetics

All initial datasets contained 42 taxa (36 ingroup Arsenurinae and 6 outgroup Saturniidae lineages). We evaluated how museum specimens performed in sequencing (i.e., papered or pinned), and did not observe any qualitatively significant differences in recovery of genetic data (Supp. Table 1, Supp. Text 2). Importantly, our datasets do not possess GC-bias (Bossert et al. 2017), with 41.7% GC content. The sequence information found within the ‘Pr’ dataset comprised 185,007 bp, of which 55,209 were informative sites, and a - 1479454.703 log-likelihood score for the consensus tree. The sequence information found within the ‘AA’ dataset comprised 60,425 sites, of which 4,709 were informative sites, and a -294666.796 log-likelihood score for the consensus tree. The sequence information found within the ‘Pr+Fl’ dataset comprised 267,958bp, of which were 93,428 informative sites, and a -2273571.993 log-likelihood score for the consensus tree. The sequence information found within the ‘Fl’ dataset comprised 80,276 bp, of which 35,647 were informative sites, and a - 817959.106 log-likelihood score for the consensus tree. See Supplemental Trees for these material.

In the initial phylogenetic inferences, we only see two recurring outcomes: 1) All genera are monophyletic entities and well-supported; and 2) *Almeidaia* – the only representative of tribe Almeidaiini – is always sister to the rest of the Arsenurinae, a clade forming the Arsenurini tribe. Generic relationships within this latter tribe are rather unresolved, except for the sister group relationship between *Dysdaemonia* and *Titaea*. We often found *Copiopteryx* and *Rhescyntis* as sister lineages, with *Paradaemonia* as the sister lineage to those two; the ‘AA’ and ‘Pr’ inferences identify the same three lineages as a monophyletic group, but with differing lineages as the sister to the other two (see Supplemental Trees). From the ‘AA’ and ‘Fl’ datasets, we recovered a “well-supported” placement of *Loxolomia* as the sister to the rest of the Arsenurini, whereas in the ‘Pr’, ‘Pr+Fl’, and ASTRAL datasets *Arsenura* is found in that position. The lineages whose topological locations change depending on the dataset are *Caio, Grammopelta*, and *Loxolomia*. In the ASTRAL tree, these three genera were never found as sister to a single other genus, but always form independent lineages branching from the tree stem (see Supplemental Trees). In the ‘AA’ tree, *Caio* is sister to *Arsenura*, and *Grammopelta* is sister to *Copiopteryx, Paradaemonia* and *Rhescyntis*. In the ‘Pr’ tree, *Caio* is sister to *Titaea* and *Dysdaemonia*, and *Grammopelta* and *Loxolomia* are sister lineages. In the ‘Fl’ tree, *Caio* is sister to *Paradaemonia*, and *Grammopelta* is sister to *Copiopteryx* and *Rhescyntis* (see Supplemental Trees). One additional common outcome is that across all trees there are very short branches and low support values associated with the early branching of the Arsenurini tribe. In particular, the ASTRAL tree, with its branch lengths in coalescent units, starkly highlights the lack of signal at this depth (see Supplemental Trees).

One of the most vexing problems in phylogenetic inference occurs when a region of a tree possesses short branches and no “support” for relationships (i.e., a polytomy). In our initial inferences, we consistently see this “unfortunate” pattern where a large polytomy (if nodes with low support are collapsed) occurs along the backbone of the Arsenurini (see Fig. 1 and Supp. Trees). Consequently, we aimed to determine whether hidden phylogenetic signal could be found in the data. First, we searched for “rogue” taxa that were potentially negatively influencing topological differences and support. Only one taxon was identified as a potential “rogue taxon”, *Grammopelta lineata* – a monotypic genus. However, *Grammopelta* was not seen as an exceptionally divergent or problematic taxon. The leaf stability index (lsDif) is a measure of the stability of a node in a set of trees, based on quartet frequencies (Thorley and Wilkinson 1999), with values ranging between 0 (unstable) and 1 (stable), but the *Grammopelta* leaf stability index, lsDif = 0.722367, was only slightly lower than those of other well supported taxa - *Dysdaemonia* and *Titaea*, both with an lsDif = 0.775075. An alternative statistic, the taxonomic instability index, measures the stability of a node in a set of trees based on unweighted patristic distances (Maddison and Maddison 2011), where the higher the annotation value, the more unstable the taxon, but the *Grammopelta* taxonomic instability index (87511.46) was again only slightly higher than other well supported taxa (e.g., *Dysdaemonia* sister to *Titaea* = 76089.91). Nevertheless, we decided to remove the taxon and rerun the phylogeny inference to investigate how topologies or support changed.

After removing *Grammopelta*, the sequence information found within the ‘AA minus *Grammopelta*’ dataset comprised 60,425 sites, of which 4,670 were informative sites, and a - 291394.450 log-likelihood score for the consensus tree. The sequence information found within the ‘Pr+Fl minus *Grammopelta*’ dataset comprised 267,958 bp, of which 92,647 were informative sites, and a -2214908.564 log-likelihood score for the consensus tree. When we compare the topological outcomes, we see that removing *Grammopelta* generally increases “traditional” support values, placing *Paradaemonia* and *Caio* as sister lineages, and provides good support for a clade comprising these two lineages as sister to *Copiopteryx* + *Rhescyntis. Titaea* and *Dysdaemonia* are sister lineages, with *Loxolomia* the sister to those two, with good support for these three lineages as sister to the previously mentioned four, and *Arsenura* and *Almeidaia* sister to the rest of the Arsenurinae. When included, *Grammopelta* is sister to *Caio*, though not well supported, and this clade is sister to ((*Rhescyntis* and *Copiopteryx*), (*Paradaemonia*, (*Titaea* and *Dysdaemonia*))), with *Loxolomia* sister to this group, and *Arsenura* and *Almeidaia* sister to the rest of the Arsenurinae (see Supplemental Trees).

We also investigated additional modes of topological support. One alternative measure of node support is to look at concordance factors, in particular, gCF values - the percentage of “decisive” gene trees containing a particular branch of interest. Using the ‘Pr+Fl’ and ASTRAL phylogenies (see Supp. Trees and Fig. 1 for these values on the preferred tree), we see that the underlying data contain much discordance – only 17 nodes within the Arsenurinae possess a gCF ≥50 and most of those correspond to nodes that define genera as well as intrageneric nodes (see Supplemental Trees), outcomes we already felt comfortable with. For example, the individual genera within the Arsenurinae are robustly supported as monophyletic; all are ≥50 gCF. We also see that seemingly undisputed nodes (e.g., the placement of *Almeidaia* as sister to the *Arsenura*, which is then sister to the rest of the Arsenurinae) have gCF values of slightly below 50 (40.6 and 49.8 respectively) – values still relatively high when compared with other nodes. Unfortunately, again we see that there is no resolution within the medium depth nodes, simply a star phylogeny (if “poorly” supported nodes are collapsed), and it is unknown whether this represents a hard or soft polytomy.

Another alternative measure of node support utilizes Internode Certainty (IC) scores, quartet-based measures for estimating internode certainty (Zhou et al. 2020). Importantly, IC scores were designed to quantify the level of incongruence in phylogenetic data sets, whereas traditional support values (i.e., bootstrap and posterior probability) aim to predict the correctness (congruence) of the tree topology. One important assumption underlying the QP-IC score is that the four subsets of taxa around a given internal branch are correctly resolved; because of this, we tested how IC scores (specifically, the QP-IC score - quadripartition internode certainty) would change depending on alternative placements of ambiguous taxa (e.g., *Grammopelta, Loxolomia*). According to Zhou et al. (2020), scores will fall between 1 and -1, summarizing the diversity and strength of conflicting signals into a single number. When scores approach 1 or -1, this suggests that a given internal branch is either supported (1) or contested (−1), whereas scores close to 0 indicate high levels of incongruence and a lack of phylogenetic signal. When we look at these scores on the ‘Pr+Fl’ phylogeny, we see that only 10 nodes have a QP-IC score ≥0.50, again with most of these representing genera or intrageneric relationships, whereas the other nodes possess values near 0 (Supp. Text 3). Following our taxon placement tests, we found alternative placements of *Grammopelta* and other taxa did not fundamentally change the pattern or strength of QP-IC score, and so only these scores are reported. For genera, we see there is good support for the reference topology (i.e., all genera are monophyletic) versus the frequency of the most prevalent alternative topology. We also find support for *Almeidaia* as sister to the rest of the Arsenurinae, but very little support for any between-genera relationships, outside of *Titaea* and *Dysdaemonia*. Unfortunately, this measure of support again confirms the high levels of incongruence amongst the individual gene trees.

Likelihood mapping was used to examine the clustering of user-defined subgroups, in a hypothesis-testing framework, as well as to visualize the phylogenetic content of the AHE dataset (Strimmer and vonHaeseler 1997). By looking at the distribution of quartets, we see there is significant signal to expect a well-resolved tree, with signal split evenly among the quartets (Supp. Fig. 3). To reconstruct a better understanding of the Arsenurinae backbone, we computed the likelihood of relationships that could be constructed from possible quartets of the taxa. First, we looked at the subfamily level by testing whether Arsenurinae is more closely related to Agliinae, Salassinae, or Saturniinae (Supp. Fig. 4). We found that 100% of the quartets possess an (Arsenurinae + Agliinae) ∼ (Salassinae + Saturniinae) relationship. Second, at the generic level we find that a relationship of (a *Loxolomia*/*Grammopelta* clade + a *Titaea*/*Dysdaemonia* clade) ∼ (a *Copiopteryx*/*Rhescyntis* + a *Paradaemonia*/*Caio* clade) is found in 69.6% of investigated quartets (with the other subfamily representatives – i.e., outgroups, *Almeidaia*, and *Arsenura* ignored) and had the best likelihood score (−2358325.992) with the shortest total tree length (2.200) (Supp. Fig. 5). Additional combinations of hypotheses were tested, though only one had an equal score and tree length – a relationship of (*Almeidaia* + a *Loxolomia*/*Caio*/*Grammopelta* clade) ∼ (a *Copiopteryx*/*Rhescyntis* + a *Paradaemonia*/*Titaea*/*Dysdaemonia* clade) is found in 51.6% of quartets, whereas (*Almeidaia* + a *Copiopteryx*/*Rhescyntis* clade) ∼ (a *Loxolomia*/*Caio*/*Grammopelta* clade + a *Paradaemonia*/*Titaea*/*Dysdaemonia* clade) represents 37.9% of quartets (the other subfamily representatives and *Arsenura* were ignored) (Supp. Fig. 6).

To evaluate gene tree/species tree incongruence in an alternative framework, we computed phylogenetic networks. Networks can help visualize reticulate relationships among taxa (i.e., due to hybridization, horizontal gene transfer, or recombination). This approach allows for comparison of the underlying phylogenetic signal because phylogenetic trees exclude all incompatible edges, whereas phylogenetic networks include them; the exclusion of an edge from a network is a powerful statement about relationships (Schliep et al. 2017). The inferred SplitsTree4 network indicates low amounts of reticulation and a large star relationship, with no intergeneric structure, except that *Loxolomia* and *Grammopelta* are closely related (Supp. Fig. 7). Unfortunately, no significant information could be gathered from the PhyloNet analyses because each time a new reticulation event was added, the likelihood score would become better, with no apparent stabilization or asymptotic behavior. Though we attempted to look at up to 14 reticulation events (i.e., equal to the total number of tips in the reduced tree), the computational resources needed (both computing power and time), resulted in this not being feasible to reasonably arrive at an answer for all 14; we tested 1 to 9 events.

Overall, to infer the most probable relationships within the Arsenurinae, we inspected the differing phylogenetic inferences, alternative support values (e.g., concordance factors, IC scores), quartets (e.g., Likelihood Mapping), and phylogenetic networks. We then inferred two unrooted constrained topologies. The constrained inference (#1) produced a consensus tree with a -2273610.971 log-likelihood score (Fig. 1). The constrained inference (#2) produced a consensus tree with a -2273675.522 log-likelihood score. We then tested whether the constrained topologies (#1 and #2) were better than the ‘Pr+Fl’ topology or each other, by using Shimodaira-Hasegawa tests under 10,000 bootstrap replicates. We find the constrained tree (#1) is not significantly better than the ‘Pr+Fl’ tree, but the constrained tree (#1) is significantly better than the constrained tree (#2). We could not statistically compare these trees with the ‘Pr+Fl minus *Grammopelta’* tree because of the missing taxon (see Supplemental Trees).

Collectively, these findings indicate *Almeidaia* is the sister lineage to all other Arsenurinae, with *Arsenura* likely the next branching lineage after *Almeidaia*. The results also indicate that within the Arsenurini *Copiopteryx* and *Rhescyntis* are sister lineages, and that *Titaea* and *Dysdaemonia* are also sister lineages. The problematic lineages (i.e., those that topologically unstable) are *Caio, Grammopelta*, and *Loxolomia*. When *Grammopelta* is removed from the inference (‘Pr+Fl minus *Grammopelta’*), *Caio* is sister to *Paradaemonia*. When *Grammopelta* is allowed to place itself in the phylogeny (constrained #2), it is inferred to be the sister lineage of a (*Copiopteryx* + *Rhescyntis*) + (*Caio* + *Paradaemonia*) clade, albeit weakly supported. In the ‘Pr+Fl’ and constrained (#1) trees, *Grammopelta* and *Loxolomia* are sister lineages, and *Caio* is sister to *Paradaemonia* (see Supplemental Trees). A synthesis of these findings provides our best estimate of the Arsenurinae phylogeny, the constrained tree (#1) (Fig. 1), and guides the trait evolution analyses (below).

### Elliptical Fourier Descriptor analysis

To examine the evolution of wing shape across the Arsenurinae, we conducted geometric morphometric analysis of 968 male specimens, mostly from natural history collections, comprising 174 species across the eight Saturniidae subfamilies (see Supp. Tables 2 & 3). Ingroup morphological sampling comprised 545 specimens corresponding to 50 species of the currently recognized 89 Arsenurinae species (see (Kitching et al. 2018); Supp. Tables 4 & 5). Across the Saturniidae, HW morphospace was visualized for all specimens at the subfamily, tribe, genus, and species level, as well as by HW shape category descriptor. Within the Arsenurinae (i.e., ingroup), morphospace was visualized at the genus and species level, as well as by HW shape category descriptor (Figs. 2 & 3). FW morphospace was plotted and visualized for the same categories, with particular focus on the HW shape category, to visualize potential trait relationships (Figs. 4 & 5). Background shapes in the morphospace plots are hypothetical shape space based on the specimens being analyzed.

**Figure 3.**
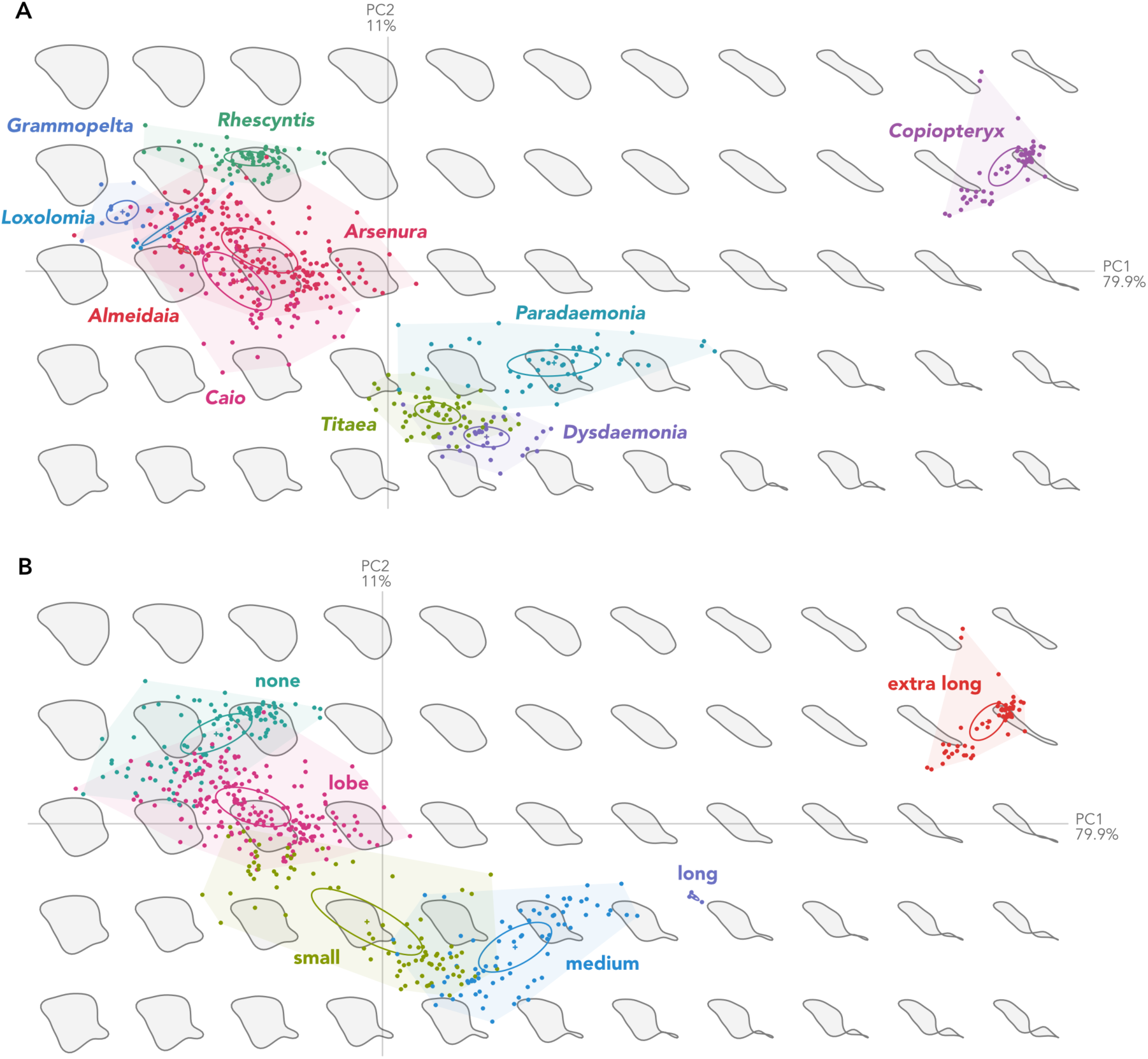
Geometric morphometric analysis of hindwing (HW) shape reveals changes in morphospace across the Arsenurinae genera. Principal components (PC1∼PC2) are plotted to visualize hindwing morphospace by A) Genus; and B) Hindwing (HW) shape morphospace plotting the hindwing tail categories “none”, “lobe”, “small”, “medium”, “long”, and “extra long”. Dots represent individual specimens in the analysis. Hypothetical shape approximations are plotted in the background to aid in visualizing shape change.

**Figure 4.**
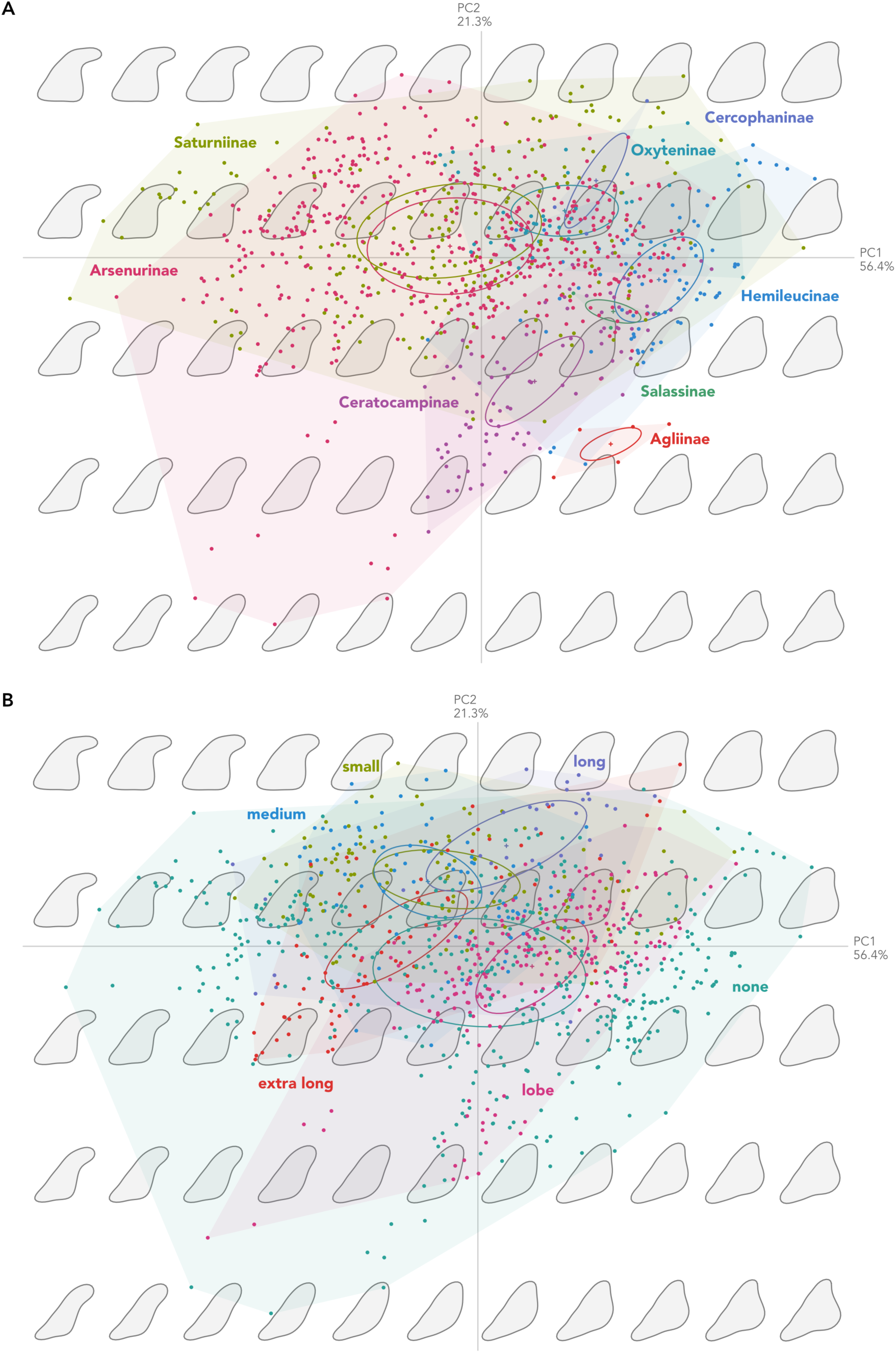
Geometric morphometric analysis of forewing (FW) shape reveals changes in morphospace across the Saturniidae subfamilies. Principal components (PC1∼PC2) are plotted to visualize forewing morphospace by A) Subfamily, showing Saturniinae and Arsenurinae have convergently evolved into the same morphospace; and B) Forewing (FW) shape morphospace plotting the hindwing tail categories “none”, “lobe”, “small”, “medium”, “long”, and “extra long” overlaid on FW shape. Dots represent individual specimens in the analysis. Hypothetical shape approximations are plotted in the background to aid in visualizing shape change.

**Figure 5.**
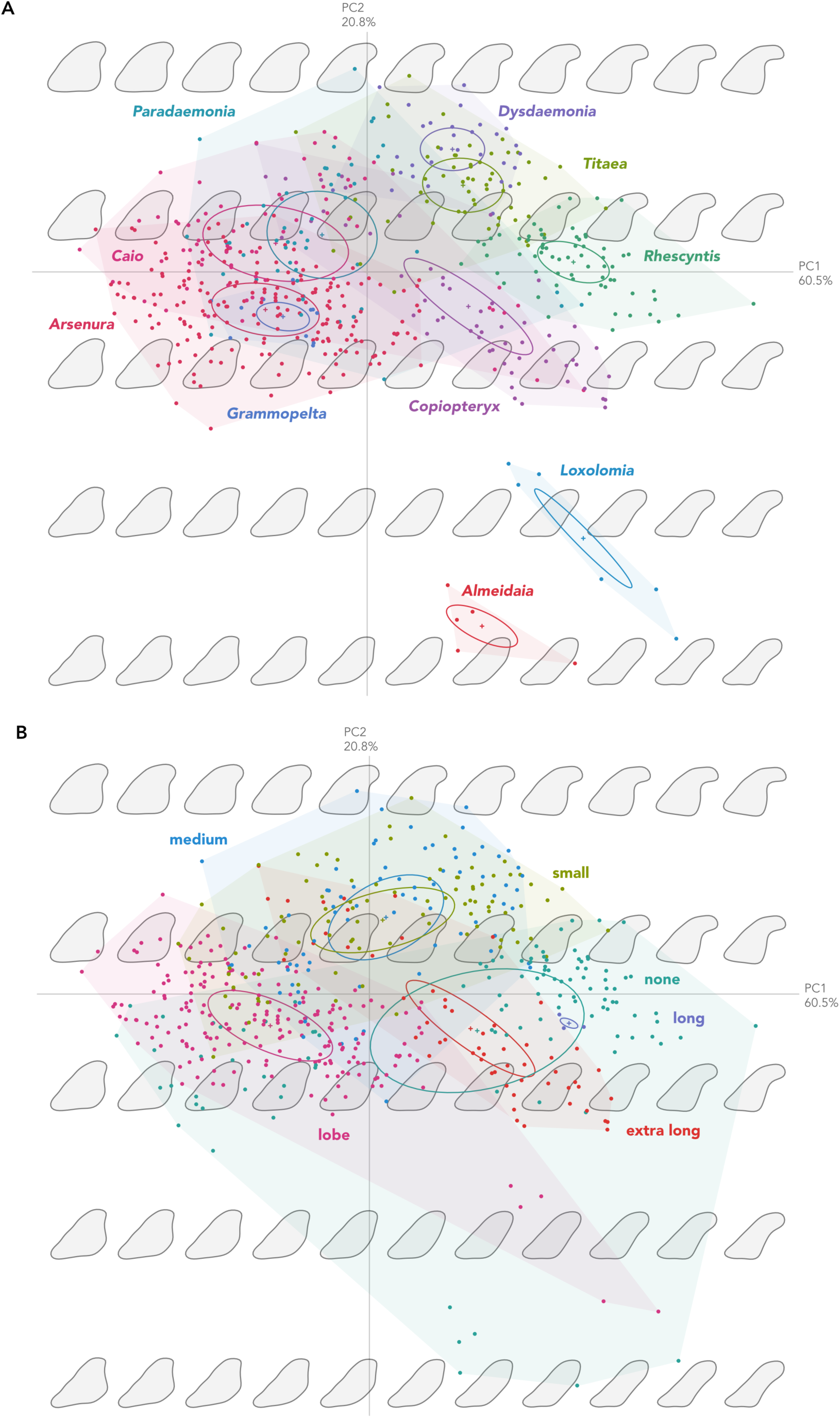
Geometric morphometric analysis of forewing (FW) shape reveals changes in morphospace across the Arsenurinae genera. Principal components (PC1∼PC2) are plotted to visualize forewing morphospace by A) genus; and B) Forewing (FW) shape morphospace plotting the hindwing tail categories “none”, “lobe”, “small”, “medium”, “long”, and “extra long” overlaid on FW shape. Dots represent individual specimens in the analysis. Hypothetical shape approximations are plotted in the background to aid in visualizing shape change.

When HW morphospace is plotted, we observe a crescent shape of morphospace that is filled by all sampled genera across the family (Fig. 2). If we classify specimens by subfamily, we see that the Arsenurinae and Saturniinae generally occupy the same morphological space, independently, as found by Rubin and Hamilton et al. (2018), suggesting there are likely evolutionary forces (e.g. bat predation and genomic constraints) driving lineages into this similar shape space (Fig. 2a). We see there is overlap in occupied shape space between HW shape categories ‘none’ and ‘lobe’, ‘lobe’ and ‘small’, ‘small’ and ‘medium’, but ‘long’ and ‘extra long’ are distinct (Fig. 2b). When we examine just the Arsenurinae (ingroup), again we see the same areas/paths of morphospace being filled, but this time it is more clear that there are three distinct shape spaces – with no overlap between *Copiopteryx* (1), *Dysdaemonia, Paradaemonia*, and *Titaea* (2), and *Almeidaia, Arsenura, Caio, Grammopelta, Loxolomia*, and *Rhescyntis* (3) (Fig. 3a). When we look to see if there is overlap in occupied shape space among the HW shape categories in the Arsenurinae, we see overlap between ‘none’ and ‘lobe’, ‘lobe’ and ‘small’, ‘small’ and ‘medium’, but ‘long’ and ‘extra long’ are distinct (Fig. 3b). Additional Arsenurinae (within genera) plots can be found in the Supplemental Information.

When we examine FW shape, we again see that the Saturniinae and Arsenurinae overlap in morphospace, though this overlap also occurs more broadly across the family (Fig. 4a). When we just examine the Arsenurinae, we see similar overlap and variation, but certain genera are distinct (*Almeidaia, Dysdaemonia, Loxolomia, Rhescyntis*, and *Titaea*) from others (*Arsenura, Caio, Grammopelta*, and *Paradaemonia*) (Fig. 5a). If we look at whether there is a potential link between FW shape and HW shape, patterns are hard to perceive due to a large amount of variation and overlap (Fig. 4b). When evaluating only the Arsenurinae, an observable pattern becomes clearer between FW shape and HW shape (Fig. 5b); those with ‘long’ or ‘extra long’ hindwings are similar, as well as ‘small’ and ‘medium’, suggesting there may be physiological/developmental “rules” to wing shape. And lastly, from a taxonomic standpoint an interesting outcome was revealed – EFD wing shape morphometrics can be used for species identification/delimitation, at least when investigating male specimens (e.g., within each genus, there are significant differences between certain species’ wing shape morphospace, in particular *Caio* and *Paradaemonia*; see the additional Arsenurinae genera plots found in the Supplemental Information).

### Trait Evolution

The main aim of this research was to determine whether there are evolutionary relationships between body size (measured as FW length - a proxy for body size), wing shape (quantified as principal components), and the presence of tails (Table 1; see Fig. 6) in the Arsenurinae, in the hope of testing for these patterns across the Saturniidae in the future. The first four principal components were used in the trait evolution analyses, with the first two the most informative (i.e., PC1 and PC2 explained ≥75% of the variation in both FW and HW shape). Before investigating the evolution of these traits across the Arsenurinae, we evaluated whether there was significant phylogenetic signal in our data (Pagel’s lambda = 0.974). A likelihood ratio test suggested there was significant signal in the phylogenetic residuals (p = 2.186e-22) (i.e., close relatives are more similar than random pairs of species; Revell 2010).

**Figure 6.**
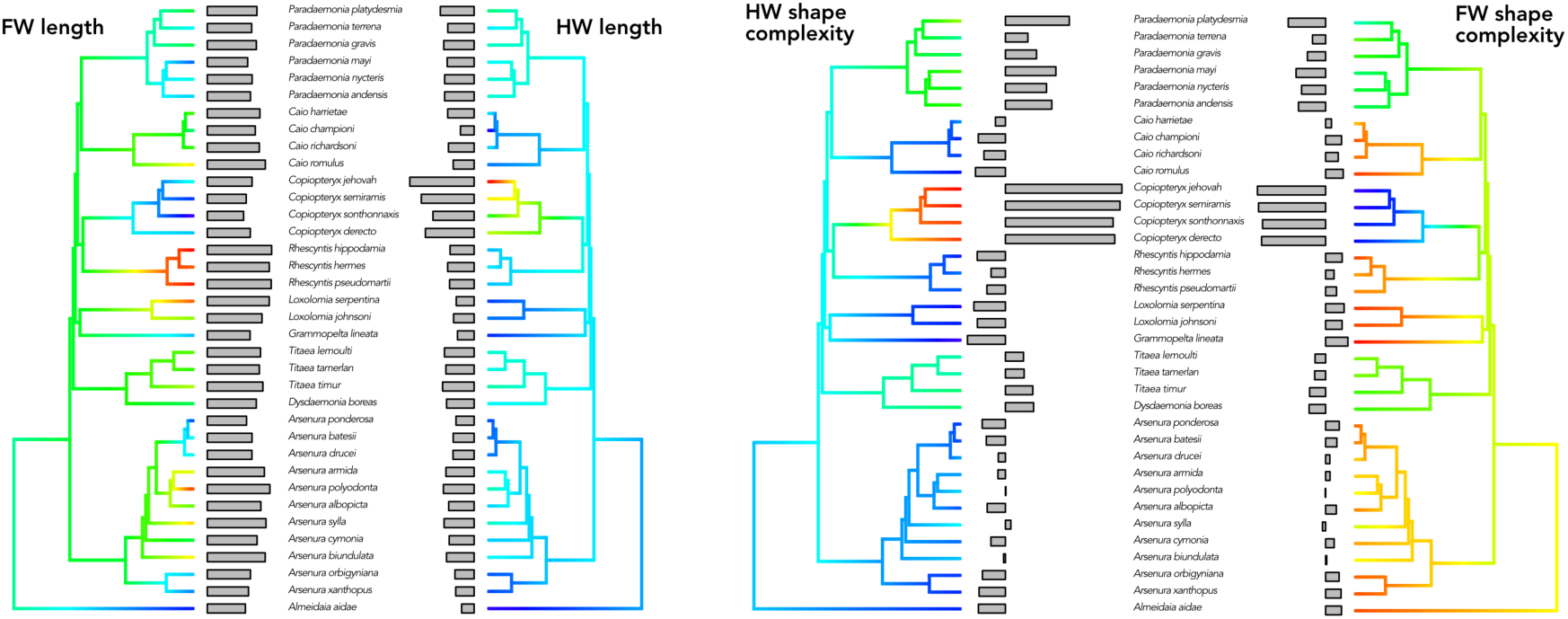
Trait evolution across the Arsenurinae phylogeny to visually inspect whether there are evolutionary relationships between body size (Left; measured as FW length - a proxy for body size), HW length (middle Left), HW shape complexity (middle Right; as principal components); and FW shape complexity (Right; as principal components).

To answer our questions relating to wing shape and body size tradeoffs, we used phylogenetic generalized least squares regressions (PGLS) and phylogenetic ANOVAs. We found that HW length (HW_L) is a predictor of body size (FW_L) (i.e., FW_L ∼ HW_L; p = 0.001). When we ask if body size determines the length of HW a moth lineage will have, we find that body size does not predict the length of the hindwing (i.e. HW_L ∼ FW_L; p = 0.1131). When we ask whether the length of the HW determines forewing shape, we find that HW length (HW_L) is a predictor of FW shape complexity (PC1 of FW shape) (i.e., PC1 ∼ HW_L; p = 2.2e-16). And lastly, we do not find that FW shape complexity predicts HW length (i.e., HW_L ∼ FW_PC1; p = 0.8423). See Table 1 for questions tested and the outcomes. See Figure 6 for shape changes across the phylogeny. To investigate the effects of HW shape category (i.e., whether or not there is a HW tail and if so what kind) on body size and FW shape we employed phylogenetic ANOVAs. We find there was no significant effect of HW shape category on body size (p = 0.502), but there was a significant effect of HW shape category on FW shape complexity (for both FW_PC1 and FW_PC2; p = 0.001). See Supp. Figures 8 and 9 for the changes in principal component morphospace across the HW and FW shapes.

## Discussion

For biologists to understand the evolution of traits, measured variables must be investigated within a solid phylogenetic framework. Phylogenomics has provided systematists with the capability to gather vast amounts of genomic sequence data in the hope of establishing those robust foundations from which evolutionary hypotheses can be tested. This molecular framework can then be integrated with other data (e.g., ecological, behavioral, morphology) to better understand how divergent forms and functions have evolved. To produce the first molecular phylogeny of the subfamily Arsenurinae, we employed Anchored Hybrid Enrichment (AHE) targeted-sequencing phylogenomics ((Lemmon et al. 2012), (Breinholt et al. 2018)). Within this phylogenetic context, we investigated the evolution of wing shape across the subfamily by integrating geometric morphometrics from natural history collection specimens to ask whether there were evolutionary trade-offs between body size, wing shape, and the presence of hindwing tails.

### Phylogeny

One (hopeful) outcome of phylogenomics is that a tree will be inferred with well-resolved nodes so that robust evolutionary inferences can be made. But even in the “age of phylogenomics” there are areas of the Tree of Life where it seems that no matter the amount of data thrown at a problem, robust resolution remains elusive. While “unfortunate”, these parts of the Tree of Life are also often some of the most interesting from an evolutionary standpoint. The Neotropical subfamily Arsenurinae appears to be one of these cases. Following initial phylogenetic inference, a large polytomy remained unresolved in the backbone of the Arsenurinae phylogeny, although all genera are monophyletic and well-supported (see Fig. 1 and Supp. Trees). One important aim was then to determine if there was hidden phylogenetic signal not being detected in traditional phylogenetic inference, and if so, where and how could it be used to piece together the evolutionary history of the group into a meaningful story.

To investigate whether hidden “support” could be found within this polytomy (i.e., a putative rapid diversification), we inspected tree topology (both supermatrix and species tree), rogue taxon analysis, alternative nodal support values (e.g., concordance factors, IC scores), quartets (e.g., Likelihood Mapping), network analyses, and constrained topology tests to arrive at a “most probable” relationship within the Arsenurinae (Fig. 1). Collectively, these analyses strongly indicate that *Almeidaia* is the sister lineage to all other Arsenurinae (the Arsenurini tribe), and that *Arsenura* is likely the next branching lineage. The results also indicate that *Copiopteryx* and *Rhescyntis* are sister lineages, as well as *Dysdaemonia* and *Titaea*. The phylogenetic network analysis indicates that *Grammopelta* and *Loxolomia* are closely related sister lineages, a finding also seen in Peigler (1993). When we compare our results to those hypothesized past relationships (Supp. Fig. 1; (Michener 1952), (Peigler 1993), (De Camargo et al. 2009)), we find little consistency (Fig. 1), except for the placement of *Almeidaia* and the sister relationship between *Dysdaemonia* and *Titaea*.

The ASTRAL tree is particularly informative because its topological structure and branch lengths (represented in coalescent units) reveal a likely diversification event in this group’s evolutionary history – an event that led to the evolution of the major genera. Though the conflicting signal that we discovered produces a polytomy, this could represent an important evolutionary event, depending on whether it is a “hard” or “soft” polytomy. Hard polytomies represent a hypothesis that a common ancestral lineage speciated into multiple lineages at the “exact” same time, although most inferred polytomies are thought to be soft – meaning that they would be resolved if data of higher quality were available. While our data are comprehensive, they are not representative of the entire genome. But interestingly, an independent phylogenomic dataset (co-author RR, in prep) using ultraconserved elements (UCE) also faces the exact same problem, supporting the hypothesis of a hard polytomy.

There are multiple additional reasons for why we find this polytomy, for example the phylogenetically-informative sites may be scattered in the “noisy” genes, such that individual gene trees do not possess much useful information. This would help answer why gCF values are low; our ability to resolve single locus gene trees is limited due to their short length (i.e., lack of molecular evolutionary signal). For example, the gCF value can be as low as 0% if no single gene tree contains a branch present in the reference tree. This is an important point because relationships with short branch lengths are hard to resolve, particularly with short loci – a common feature of target capture phylogenomic data, indicating that many of our loci contain genuinely conflicting signal. Lastly, all but two genera in the subfamily have fewer than 10 described species (*Grammopelta* = 1 species; *Almeidaia* = 2; *Loxolomia* = 3; *Rhescyntis* and *Titaea* = 5; *Dysdaemonia* = 6; *Caio* and *Copiopteryx* = 7), whereas *Paradaemonia* and *Arsenura* have 20 and 34 species, respectively. It is possible that many lineages have gone extinct, but at this time it is unknown whether this disparity was due to an elevated extinction rate in some lineages, a mass extinction event, a generic-level burst of diversification within this relictual lineage, or simply that a lack of taxonomic effort has not yet discovered the full extant diversity in this subfamily. Based on our previous knowledge of the bat-moth arms race, we propose a burst of diversification occurred in the Arsenurinae (as evidenced by the phylogeny), but at this time it is unknown what exactly drove the burst. A dated phylogeny would help us answer this question, but due to the paucity of informative Saturniidae and Bombycoidea fossils (Breinholt and Kawahara 2013) this was not undertaken. Our findings suggest this region of the Tree of Life may truly be unresolvable. But at this time, we cannot pinpoint the mechanism that allowed for this rapid split of lineages, perhaps more taxon sampling combined with biogeographical and ecological data will better answer this in the future.

### Morphology & Trait Evolution

The Arsenurinae is not a “species rich” lineage of Saturniidae, but their diverse wing shape morphospace and discovery to be a relictual lineage (Hamilton et al. 2019) can shed significant light on the evolution of the Saturniidae. The subfamily possesses a number of wing shapes that are shared with other subfamilies (see Rubin et al. 2018). For example, Arsenurinae and Saturniinae occupy the same morphological space, suggesting there are likely multiple evolutionary forces (e.g. bat predation and genomic constraints) driving these lineages into this convergent shape space (Figs. 2a & 4a). Future work will have to parse whether these forces are more heavily weighted towards genetic constraints, or whether natural selection (perhaps by bat predation) has played the larger role. We are only now beginning to answer some of the more fascinating questions in the evolution of the Saturniidae – e.g., whether tails evolved using the same path, over and over, and whether long hindwing tails gradually evolved in a progressive manner from no tails. Similar to the findings of Rubin and Hamilton et al. (2018), our results indicate that hindwing tails evolved rapidly and not transitionally.

The most ecologically elucidating aspect of this investigation was determining there were evolutionary trade-offs between body size, forewing shape, hindwing length, and the presence of tails (Table 1; see Fig. 6): 1) body size and the length of the HW covary inversely to one another, such that as HW length increases, body size decreases – interestingly, this trait (i.e., a very small body size and extra-long tails) has evolved independently in the long-tailed African saturniid genus *Eudaemonia* (Rubin et al. 2018); 2) the complexity of FW shape also covaries in opposition to the length of the HW, such that as HW length increases, FW shape becomes less complex; and 3) the type of HW shape category that a lineage possesses has an effect on FW shape, such that as lineages move through the shape states from no tail to long tail, the FW becomes less complex. These results also make intuitive sense – for example, the largest body sizes in the Arsenurinae are found in the genus *Rhescyntis*, a lineage without hindwing tails (Fig. 6) but instead possess large lobed hindwings (Fig. 1), a trait experimentally shown to reduce capture success by bats, as well as having evolved independently in the Saturniidae tribe Attacini (Rubin et al. 2018). And lastly, there are a number of saturniid lineages that are small and do not have hindwing tails. While we see that FW shape complexity does not predict HW length, those Arsenurinae possessing a long hindwing tail exhibit less complex FW shape – a finding likely important on a macroevolutionary scale, given the possible flight constraints imposed by carrying long hindwing tails (reviewed in Le Roy et al. (2019)).

There are many reasons why these tradeoffs might occur across evolutionary time. Perhaps there may be an energetic cost to building a long hindwing tail during pupal development, which might limit the resources available for other structures (such as large bodies or convoluted forewing shapes) (Shingleton et al. 2007). Alternatively, it could be that because large body size is also a corollary of being a capital breeder (i.e., needing to store resources as a larvae for future flight as adults because the lineage does not nectar feed as adults), Saturniidae lineages cannot possess a large body size if they also have long tails – potentially explaining why we do not see more species with tails across the family. Detailed kinematic analysis of diverse silkmoths in flight could provide the explanatory power, as it is possible that increasing the complexity of the hindwings imposes limitations on body size and FW shape – structures more critical to moth flight than hindwings (Jantzen and Eisner 2008). For example, based on our observations (RR pers. obs.), long-tailed saturniids (e.g., *Copiopteryx* and *Eudaemonia*) have a different flight behavior than non-tailed species – i.e., long-tailed species tend to flutter highly erratically yet surprisingly maneuverable. In contrast, genera like *Arsenura* and *Titaea* exhibit a more powerful flight, potentially due to the larger wing surface area. Future research should also look into whether less complex forewing shape creates more uniform power during flight. Additionally, butterfly fore- and hindwings appear to be subject to differing selective pressures (Owens et al. 2020) and have been linked with microhabitat (Chazot et al. 2016), which will be an interesting question to consider in future research, as members of the Arsenurinae, and Saturniidae at large, have a wide geographical spread across a variety of environments. It has also recently been seen that, in butterflies, the hindwing is important in gliding performance (Stylman et al. 2019). And lastly, research on the effects of wing damage to specific regions of the fore- and hindwings indicates that altering the shape of the leading edge of the forewing has the greatest impact on flight parameters (Le Roy et al. 2019). We suggest that more work is needed in the Saturniidae family in general to understand the kinematic effects of complex FW and HW shape.

Arsenurine moths, like other saturniids, do not possess many of the anti-bat traits that can be found in their sister lineage, the sphingids, such as ears or ultrasound producing organs. One intriguing, but as yet untested, hypothesis is that body size and FW shape are anti-bat strategies. Insectivorous bats are gape-limited predators (Santana 2016). This functionally-restricted morphology may restrict bats to attacking only “appropriately sized” prey. Thus, large-bodied moths may effectively reduce or eliminate predation events from small bat species. In fact, some of the largest moths are thought to avoid predation simply because their size prevents bats from capturing them (Roeder and Treat 1970). Small myotid bats, when offered dead, tethered moths of various sizes, have been shown to attack small moths approximately 50% more frequently than large ones (Barclay and Brigham 1994). In addition, while experimental evidence indicates that hindwing tails are a highly-effective anti-bat evolutionary strategy that has evolved multiple times within the Saturniidae, it is not yet clear whether complex forewing shape can function to thwart bat attack ((Barber et al. 2015), (Rubin et al. 2018)). Our present morphometric and phylogenetic investigation into the Arsenurinae suggests that these hindwing appendages come with a body size and forewing shape trade-off. These results provide another layer of insight into the evolutionary forces structuring wing shape in the Arsenurinae and likely the Saturniidae.

## Acknowledgements

We thank Vincent Bonhomme for his help with Momocs. We are indebted to Charles Mitter and Kim Mitter for the use of their specimens in the UMD collection, as well as Jon Heppner for the use of specimens in his collections (FLMNH) – this work would absolutely not have been done without their help. We thank those who helped collect specimens (too many to list here). Kelly Dexter assisted with DNA extractions and additional lab work; Ryan St. Laurent provided input with Arsenurinae systematics. Sammantha Epstein and the FLMNH volunteer team in the Kawahara Lab helped voucher specimens. UF undergraduates Adena Mahadai, Shaelyn McGiveron, Dominique Philoctete, and Neeka Sewnath helped with specimen digitization, as did UF researcher Geena Hill and visiting researchers James Adams and John Snyder. We acknowledge the UF HPC for providing computational support and assistance. Hamilton thanks AYK and Charles Cobb for their guidance as sponsoring scientists during the PRFB.

## Data availability

Data available from the Dryad Digital Repository: http://dx.doi.org/10.5061/dryad.[NNNN]

## Appendices

### Supplemental Figures

<u>Supplemental Figure 1</u> – The past hypothesized Arsenurinae relationships of Michener (1952) (left), Peigler (1993) (middle), and De Camargo et al. (2009) (right), reproduced for this publication.

<u>Supplemental Figure 2</u> – Illustrations for how forewing (FW) length and hindwing (HW) length measurements were made.

<u>Supplemental Figure 3</u> – Quartet likelihood mapping to inspect the phylogenetic content of the dataset. By looking at the distribution of quartets, we see that there is significant signal to expect a well-resolved tree, with signal split evenly among the quartets.

<u>Supplemental Figure 4</u> – Quartet likelihood mapping of possible relationships at the subfamily level. We tested whether Arsenurinae was more closely related to the Agliinae, Salassinae, or Saturniinae. We found that 100% of the quartets possess an (Arsenurinae + Agliinae) ∼ (Salassinae + Saturniinae) relationship.

<u>Supplemental Figure 5</u> – Quartet likelihood mapping of possible relationships at the genus level. We tested relationships by grouping lineages, according to the phylogeny, and evaluating (from a quartet standpoint) whether lineages were found more closely related to another. We find that a relationship of (a *Loxolomia*/*Grammopelta* clade + a *Titaea*/*Dysdaemonia* clade) ∼ (a *Copiopteryx*/*Rhescyntis* + a *Paradaemonia*/*Caio* clade) is found in 69.6% of investigated quartets (with the other subfamily representatives, *Almeidaia*, and *Arsenura* ignored).

<u>Supplemental Figure 6</u> – Quartet likelihood mapping of possible relationships at the genus level. We tested relationships by grouping lineages, according to the phylogeny, and evaluating (from a quartet standpoint) whether lineages were found more closely related to another. We find that a relationship of (*Almeidaia* + a *Loxolomia*/*Caio*/*Grammopelta* clade) ∼ (a *Copiopteryx*/*Rhescyntis* + a *Paradaemonia*/*Titaea*/*Dysdaemonia* clade) is found in 51.6% of quartets, whereas (*Almeidaia* + a *Copiopteryx*/*Rhescyntis* clade) ∼ (a *Loxolomia*/*Caio*/*Grammopelta* clade + a *Paradaemonia*/*Titaea*/*Dysdaemonia* clade) represents 37.9% of quartets (the other subfamily representatives and *Arsenura* were ignored). This relationships was the only other of those tested that had an equal score and tree length to Supplemental Figure 4.

<u>Supplemental Figure 7</u> – To investigate potential gene tree/species tree incongruence, we computed a phylogenetic network. The inferred SplitsTree4 network indicates low amounts of reticulation and a large star relationship, with no intergeneric structure, except that *Loxolomia* and *Grammopelta* are closely related

<u>Supplemental Figure 8</u> – Plot of changes in hindwing (HW) PC morphospace.

<u>Supplemental Figure 9</u> – Plot of changes in forewing (FW) PC morphospace.

<u>Directory “Arsenurinae_genera_morphospace_plots”</u> – HW and FW plots for all Arsenurinae genera and species investigated herein.

### Supplemental Tables

<u>Supplemental Table 1</u> – Tab-delimited file of information for specimens used in phylogenetic inference. Included are the specimen codes, taxonomy, their condition (i.e., molecular, papered, or pinned specimen), and year they were collected.

<u>Supplemental Table 2</u> – Tab-delimited file of hindwing (HW) images used to quantify the shape complexity (as principal components) of 968 male specimens comprising 174 species across the eight Saturniidae subfamilies.

<u>Supplemental Table 3</u> – Tab-delimited file of forewing (FW) images used to quantify the shape complexity (as principal components) of 968 male specimens comprising 174 species across the eight Saturniidae subfamilies.

<u>Supplemental Table 4</u> – Tab-delimited file of ingroup (i.e., the Arsenurinae) hindwing (HW) images used to quantify the shape complexity (as principal components) of 545 male specimens comprising 50 of the 89 currently recognized species.

<u>Supplemental Table 5</u> – Tab-delimited file of ingroup (i.e., the Arsenurinae) forewing (FW) images used to quantify the shape complexity (as principal components) of 545 male specimens comprising 50 of the 89 currently recognized species.

### Supplemental Text

<u>Supplemental Text 1</u> – Tab-delimited file of locus recovery for each AHE locus used in phylogenetic inference. A cutoff of ≥50% was applied to the sampled taxa (i.e., for a locus to be included in the analysis, the locus had to be recovered in at least 50% of the sampled taxa).

<u>Supplemental Text 2</u> – Tab-delimited file of taxon recovery for each AHE taxon used in the phylogenetic inference.

<u>Supplemental Text 3</u> –The file is a text file with a newick tree embedded within. QP-IC scores (quadripartition internode certainty) can be found at nodes.

### Supplemental Trees

<u>Arsenurinae.AA.noGrammopelta.pdf & .tre</u> – Newick tree file and PDF from the amino acid (AA) phylogeny without *Grammopelta*.

<u>Arsenurinae.AA.pdf & .tre</u> – Newick tree file and PDF from the amino acid (AA) phylogeny.

<u>Arsenurinae.PROBE.pdf & .tre</u> – Newick tree file and PDF from the nucleotide ‘Pr’ phylogeny.

<u>Arsenurinae.concat.constraint1.pdf & .tre</u> – Newick tree file and PDF from the constrained inference (#1) nucleotide dataset ‘Pr+Fl’ constrained with ‘Constraint_tree_01’.

<u>Arsenurinae.concat.constraint2.pdf & .tre</u> – Newick tree file and PDF from the constrained inference (#2) nucleotide dataset ‘Pr+Fl’ constrained with ‘Constraint_tree_02’.

<u>Arsenurinae.concat.noGrammopelta.pdf & .tre</u> – Newick tree file and PDF from the nucleotide ‘Pr+Fl’ phylogeny without *Grammopelta*.

<u>Arsenurinae.concat.pdf & .tre</u> – Newick tree file and PDF from the nucleotide ‘Pr+Fl’ phylogeny.

<u>Arsenurinae.flanking.pdf & .tre</u> – Newick tree file and PDF from the nucleotide ‘Fl’ phylogeny.

<u>Constraint_tree_01.pdf & .tre</u> – Newick tree file and PDF of the ‘constrained inference (#1)’ phylogeny.

<u>Constraint_tree_02.pdf & .tre</u> – Newick tree file and PDF of the ‘constrained inference (#2)’ phylogeny.

<u>concordance.factor.ASTRAL.tree.pdf & .tre</u> – Newick tree file and PDF from the ASTRAL inference, with concordance factors placed at nodes.

<u>concordance.factor.concat.tree.pdf & .tre</u> – Newick tree file and PDF from the constrained inference (#1) nucleotide dataset, with concordance factors placed at nodes.

